# ApoA-I Protects Pancreatic β-cells from Cholesterol-induced Mitochondrial Damage and Restores their Ability to Secrete Insulin

**DOI:** 10.1101/2023.04.03.535492

**Authors:** Bikash Manandhar, Elvis Pandzic, Nandan Deshpande, Sing-Young Chen, Valerie Wasinger, Maaike Kockx, Elias Glaros, Kwok Leung Ong, Shane R. Thomas, Marc R. Wilkins, Renee M. Whan, Blake J. Cochran, Kerry-Anne Rye

## Abstract

**Objective:** High cholesterol levels in pancreatic β-cells cause oxidative stress and decrease insulin secretion. β-cells can internalize apolipoprotein (apo) A-I, which increases insulin secretion. This study asks whether internalization of apoA-I improves β-cell insulin secretion by reducing oxidative stress.

**Approach:** Ins-1E cells were cholesterol-loaded by incubation with cholesterol-methyl-β-cyclodextrin. Insulin secretion in the presence of 2.8 or 25 mM glucose was quantified by radioimmunoassay. Internalization of fluorescently labelled apoA-I by β-cells was monitored by flow cytometry. The effects of apoA-I internalization on β-cell gene expression was evaluated by RNA sequencing. ApoA-I binding partners on the β-cell surface were identified by mass spectrometry. Mitochondrial oxidative stress was quantified in β-cells and isolated islets with MitoSOX and confocal microscopy.

**Results:** An F_1_-ATPase β-subunit on the β-cell surface was identified as the main apoA-I binding partner. β-cell internalization of apoA-I was time-, concentration-, temperature-, cholesterol- and F_1_-ATPase β-subunit-dependent. β-cells with internalized apoA-I (apoA-I^+^ cells) had higher cholesterol and cell surface F_1_-ATPase β-subunit levels than β-cells without internalized apoA-I (apoA-I^−^ cells). The internalized apoA-I co-localized with mitochondria and was associated with reduced oxidative stress and increased insulin secretion. The ATPase inhibitory factor 1, IF_1_, attenuated apoA-I internalization and increased oxidative stress in Ins-1E β-cells and isolated mouse islets. Differentially expressed genes in apoA-I^+^ and apoA-I^−^ Ins-1E cells were related to protein synthesis, the unfolded protein response, insulin secretion and mitochondrial function.

**Conclusions:** These results establish that β-cells are functionally heterogeneous and apoA-I restores insulin secretion in β-cells with elevated cholesterol levels by improving mitochondrial redox balance.

## INTRODUCTION

Diabetes mellitus is a chronic condition that is characterized by elevated blood glucose levels due to inadequate insulin production and/or ineffective utilization of insulin. Diabetes is classified into two main types. Type 1 diabetes is an autoimmune disease in which pancreatic β-cells are selectively destroyed by auto-activated T cells, leading to reduced β-cell mass and decreased insulin secretion in response to glucose. Type 2 diabetes, which accounts for ∼90% of all diabetes, is caused by ineffective use of insulin (insulin resistance). β-cells compensate for insulin resistance by producing and secreting extra insulin. However, this ultimately causes β-cell exhaustion, progressively reduces β-cell mass and drives progression from pre-diabetes to overt diabetes.

Cholesterol accumulation in β-cells also impairs insulin secretion^1–3^ by reducing insulin granule exocytosis.^4, 5^ It also causes mitochondrial dysfunction^6^ and oxidative and endoplasmic reticulum (ER) stress.^7, 8^ Importantly, the deleterious effect of high β-cell cholesterol levels on insulin secretion can be reversed by depleting the cells of excess cholesterol with an extracellular cholesterol acceptor such as methyl-ý-cyclodextrin.^2^

ApoA-I, the main apolipoprotein constituent of high density lipoproteins (HDLs), improves glucose tolerance in high-fat fed C57BL6 mice by increasing β-cell insulin secretion and, in the case of insulin resistant *db/db* mice, increasing glucose uptake into skeletal muscle.^9, 10^ ApoA-I and HDLs are the main acceptors of cholesterol that effluxes from cell membranes in a process that is dependent on the ATP-binding cassette transporters ABCA1 and ABCG1.^11^ This suggests that apoA-I and HDLs may improve glucose tolerance in mice with diabetes by reducing β-cell cholesterol levels and increasing insulin secretion. However, we have found that apoA-I increases glucose-stimulated insulin secretion (GSIS) in β-cells and isolated rodent islets without altering cellular cholesterol levels.^12, 13^ For example, apoA-I treatment of mice with increased islet cholesterol levels and impaired insulin secretion due to conditional β-cell deletion of ABCA1 and ABCG1 markedly improves glycaemic control and GSIS.^14, 15^ This is consistent with apoA-I improving β-cell function independent of cholesterol levels.

It has recently been shown that rat Ins-1E insulinoma cells internalize apoA-I by a mechanism that involves translocation of the transcription factor pancreatic and duodenal homeobox 1(Pdx1) to the nucleus, increased processing of proinsulin into insulin and increasing the number of insulin granules docked at the β-cell surface.^16^ While this has provided some insight into how apoA-I increases insulin secretion without reducing β-cell cholesterol levels, the underlying mechanism remains unclear.

This question is addressed in the present study, which establishes that apoA-I internalization is increased in Ins-1E cells with elevated cholesterol levels and impaired GSIS, and that the internalized apoA-I colocalises with mitochondria where it lowers oxidative stress and increases insulin secretion.

## METHODS

### Purification of lipid free apoA-I and labelling with Alexa Fluor (AF) 488

Human apoA-I (CSL Behring, King of Prussia, PA) was purified on a Q-Sepharose Fast-Flow column attached to an ÄKTA™ FPLC system.^17^ Purity (>95%) was confirmed by SDS-PAGE and staining with Coomassie Brilliant Blue G-250 (Bio-Rad Laboratories, Hercules, CA).

ApoA-I (2–5 mg/mL) was dissolved in sodium bicarbonate (0.1 M), added to AF488 (Thermo Fisher Scientific, Waltham, MA) and maintained at room temperature for 1 h, with gentle mixing every 15 min. Unconjugated/free dye was removed by centrifugation at 1100 *g* for 5 min with a resin supplied by the manufacturer. Protein concentrations and labelling efficiency were determined using a Nanodrop 1000 spectrophotometer (Thermo Fisher Scientific). AF488-labelled apoA-I was used at a final concentration of 0.1 mg/mL, unless stated otherwise.

### Glucose-stimulated insulin secretion (GSIS)

Ins-1E cells were loaded with cholesterol by incubation with cholesterol-methyl-β-cyclodextrin (5 mM), washed with KRBH buffer and incubated without or with apoA-I (final concentration 1 mg/mL) for 2 h under basal (2.8 mM) or high (25 mM) glucose conditions. The medium was then collected, and the cells were lysed (0.2 M NaOH). The protein concentration of the cell lysates was measured using the BCA assay and insulin levels were quantified by radioimmunoassay (rat insulin RIA kit, Merck, Darmstadt, Germany).

### Cholesterol loading of Ins-1E cells

Ins-1E rat insulinoma cells were loaded with cholesterol by incubation with cholesterol-methyl-β-cyclodextrin (final concentration 5 mM). See Supplemental Methods for details.

### Quantification of mitochondrial reactive oxygen species (ROS)

The ability of apoA-I to inhibit ROS formation in cholesterol-loaded Ins-1E cells was quantified by seeding cells onto a black, transparent bottom 96-well plate. At 80% confluency the cells were incubated for 1 h with KRBH buffer without or with 5 mM cholesterol. After washing with KRBH buffer, the cells were incubated for 1 h in the absence or presence of apoA-I (final concentration 0.1-1 mg/mL), then incubated for 10 min with MitoSOX (5 µM, Thermo Fisher Scientific). ROS formation was quantified as fluorescence intensity (excitation wavelength 510 nm, emission wavelength 580 nm) using a Clariostar plate reader (Molecular Devices, San Jose, CA). The cells were then lysed, and the protein concentration was measured by BCA assay. Fluorescence intensity was normalised to protein concentration.

### Regulation of apoA-I internalization in Ins-1E cells

The concentration-dependent internalization of apoA-I by Ins-1E cells was evaluated by incubation for 1 h at 37 °C in KRBH buffer in the absence or presence of unlabelled and AF488-labelled apoA-I under basal (2.8 mM) glucose conditions (final total apoA-I concentration 0.1-1.0 mg/mL). AF488-labelled apoA-I was included in the incubations at one-tenth the concentration of unlabelled apoA-I.

ApoA-I internalization under basal (2.8 mM) and high (25 mM) glucose concentrations was evaluated by incubating Ins-1E cells at 37 °C for 1 h in KRBH buffer in the absence or presence of unlabelled apoA-I (final concentration 0.9 mg/mL) and AF488-labelled apoA-I (final concentration 0.1 mg/mL).

Time-dependent apoA-I internalization was evaluated by incubating Ins-1E cells for up to 2 h at 37 °C in KRBH buffer in the absence or presence of unlabelled apoA-I (final concentration 0.9 mg/mL) and AF488-labelled apoA-I (final concentration 0.1 mg/mL) under basal (2.8 mM) glucose conditions. AF488 fluorescence was determined at 5, 10, 15, 30, 60 and 120 min.

Temperature-dependent apoA-I internalization was evaluated by incubating Ins-1E cells for 1 h at 4 °C or 37 °C in KRBH buffer in the absence or presence of unlabelled apoA-I (final concentration 0.9 mg/mL) and AF488-labelled apoA-I (final concentration 0.1 mg/mL) under basal (2.8 mM) glucose conditions.

The impact of cholesterol loading on apoA-I internalization was evaluated by pre-incubating Ins-1E cells for 1 h in the absence or presence of cholesterol (final concentration 5 mM). The cholesterol-loaded cells were then incubated at 37 °C for 1 h in the absence or presence of unlabelled apoA-I (0.9 mg/mL) and AF488-labelled apoA-I (0.1 mg/mL) under basal (2.8 mM) glucose conditions.

When the incubations were complete, the cells were washed with ice-cold PBS (x2), harvested with PBS/EDTA-Na_2_ (2 mM) and pelleted (300 *g*, 4 °C, 3 min). The pellets were washed twice and cell surface AF488 fluorescence was quenched by incubation for 30 min at 4 °C with rabbit anti-AF488 IgG (4 µg/mL, Thermo Fisher Scientific). The difference between the signals from the cells incubated in the absence and presence of anti-AF488 IgG corresponded to apoA-I bound to the cell surface.

Live cells were identified by staining with propidium iodide (PI) (Merck). Fluorescence was measured using a FACSCanto II flow cytometer (BD Biosciences, Franklin Lakes, NJ) and analysed using FlowJo (BD Biosciences). The gating strategy is shown in Supplementary Fig 1.

**Fig 1.**
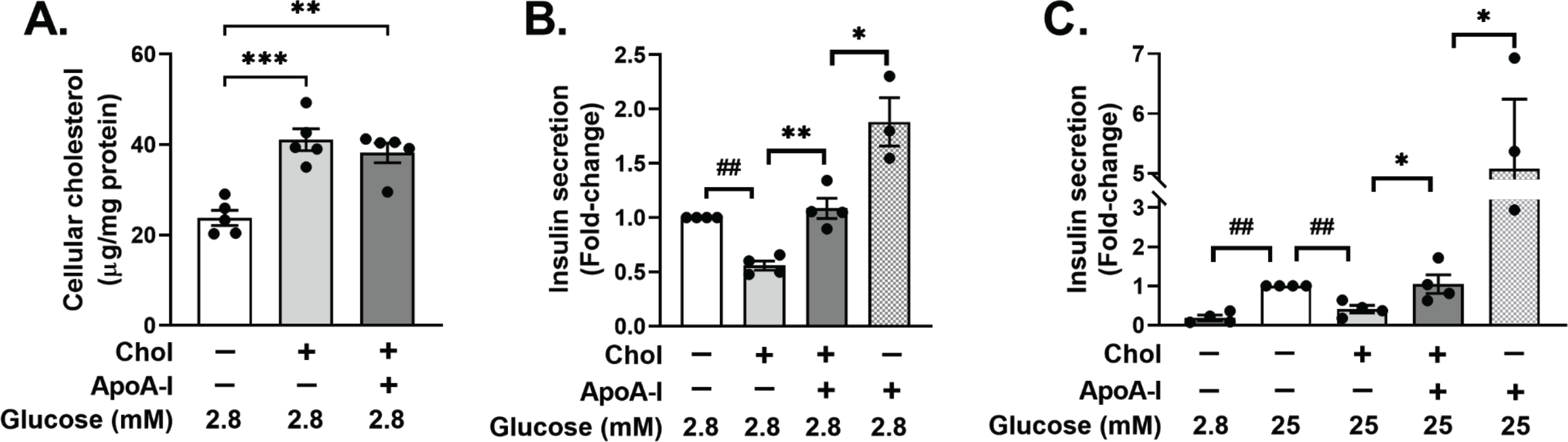
ApoA-I increases GSIS in non-cholesterol-loaded and cholesterol-loaded Ins-1E cells. Ins-1E cells were incubated for 1 h at 37 °C in the absence or presence of cholesterol-β-methyl cyclodextrin (5 mM cholesterol), washed with PBS then incubated for 2 h without or with apoA-I (final concentration 1 mg/mL). **Panel A:** Cells were incubated at 37 °C for 1 h in the presence and absence of cholesterol-β-methyl cyclodextrin. Cholesterol levels were measured by HPLC and normalised to cell protein concentration. Non-cholesterol-loaded and cholesterol-loaded cells were incubated in the presence and absence of apoA-I under basal (2.8 mM glucose, **Panel B**) and high (25 mM glucose, **Panel C**) glucose conditions. Insulin levels in the medium were quantified by RIA, normalised to cell protein concentration and expressed as fold-change relative to control. The values represent mean±SEM of 4-5 independent experiments, with 3-4 replicate samples for each independent experiment. Panel A was analyzed via 1-way ANOVA, followed by Tukey’s multiple comparison test. Panels B-C were analyzed by 1-sample t-test or unpaired, 2-tailed t-test, as appropriate. *p<0.05, **p<0.01, ***p<0.0001, ^##^p<0.01.

### Identification of the F_1_-ATPase β-subunit on the Ins-1E cell surface

Tosylactivated Dynabeads (Thermo Fisher Scientific) were used to identify apoA-I binding partners on the Ins-1E cell surface.^18^ ApoA-I (100 µg) was dissolved in borate buffer (0.1 M, pH 9.5) containing ammonium sulphate (1.2 M) and incubated at 37 °C overnight with Dynabeads (5 mg, 165 µL). The beads with bound apoA-I were collected using a magnet. Coupling efficiency was determined by measuring the unbound apoA-I concentration in the supernatant. The beads with bound apoA-I were resuspended in PBS (pH 7.4, 250 µL) with 0.1% (w/v) BSA and incubated (5 min, room temperature) on a rotary shaker. The beads were collected and resuspended in PBS (250 µL) with 0.5 % (w/v) BSA, then incubated for 1 h at 37 °C to reduce non-specific binding. The beads were collected and resuspended in PBS (250 µL) without BSA.

Ins-1E cells were seeded in a 12-well plate, washed with KRBH buffer and maintained at 4 °C for 1 h in KRBH buffer/0.1% (w/v) BSA (200 µL) and Dynabeads with bound apoA-I (50 µL). The supernatant was removed and the cells were washed with ice-cold PBS and harvested with PBS/EDTA-Na_2_ (2 mM). The Dynabeads were isolated with a magnet, washed with PBS (pH 7.4)/0.1% (v/v) Tween 20, then washed with PBS (pH 7.4). Non-specifically bound proteins and peptides were removed from the Dynabeads by incubation overnight at room temperature with trypsin (final concentration 0.1 mg/mL). The beads were collected using a magnet and washed (x3) with PBS. ApoA-I binding partners were eluted with 0.15% (v/v) trifluoroacetic acid, concentrated (C18 Spin Tips, Thermo Fisher) and analysed by LC– MS/MS (see Supplementary Methods).^18^

### Dependence of apoA-I internalization on cell surface F_1_-ATPase β-subunit in Ins-1E cells and primary islets

Cell surface F_1_-ATPase β-subunit levels were quantified by incubating Ins-1E cells for 1 h at 4 °C with a mouse IgG_1_ kappa antibody (isotype control (1:100), final concentration 10 µg/mL) or a mouse anti-F_1_-ATPase β-subunit antibody (3D5 clone, (1:50), final concentration 10 µg/mL, Abcam, Cambridge, UK). The cells were then incubated for 30 min at 4 °C with AF647-conjugated goat anti-mouse IgG (1:500, Thermo Fisher). Cholesterol levels were determined by incubation for 30 min at 4 °C with filipin (50 µg/mL). Live cells were identified with PI. Mean fluorescence intensity (MFI) was determined using a BD LSRFortessa X-20 flow cytometer (BD Biosciences).

Cell surface F_1_-ATPase β-subunit in Ins-1E cells was blocked by pre-incubation (37 °C, 15 min) with IF_1_ (final concentration 10 µg/mL, Creative BioMart, Shirley, NY) in KRBH/0.1 % (w/v) BSA (pH 6.4). The cells were then incubated (30 min, 37 °C) with AF488-labelled apoA-I (final concentration 0.1 mg/mL). The cells were washed with ice-cold PBS, harvested, and incubated (30 min, 4 °C) with rabbit anti-AF488 IgG (4 µg/mL, Thermo Fisher). AF488 fluorescence was quantified by flow cytometry (BD FACSCanto II flow cytometer, BD Biosciences).

Handpicked islets were incubated overnight (37 °C, 5 % CO_2_) in a humidified incubator (35 mm glass bottom fluoro dish, Cellvis, Mountain View, CA). The medium was removed, the islets were washed with KRBH, then incubated (37 °C, 15 min) in KRBH/0.1 % (w/v) BSA (pH 6.4) with or without the F_1_-ATPase β-subunit inhibitor, IF_1_ (final concentration 10 µg/mL). AF488-labelled apoA-I (0.1 mg/mL) was added, and the cells were incubated for 1 h at 37 °C. Hoechst 33342 (Thermo Fisher, 1:200 dilution (v/v) of a 10 mg/mL solution) was added and the islets were incubated for 10 min. The islets were washed with KRBH, fixed (15 min, room temperature, 10% (v/v) formalin) permeabilised then incubated overnight at 4 °C with AF647-conjugated rabbit insulin antibody (1:100, Cell Signalling). After washing, ProLong Diamond Antifade Mountant (Thermo Fisher) was added dropwise and the samples were maintained at 4 °C until imaged (Zeiss LSM 880 confocal microscope with Airyscan, Carl Zeiss Microscopy, Jena, Germany) as described in Supplementary Methods.

### Regulation of cholesterol-induced mitochondrial oxidative stress by internalized apoA-I

Ins-1E cells were seeded onto a 35 mm glass bottom fluorodish (Cellvis) and incubated (37 °C, 1 h) with KRBH buffer in the absence or presence of 5 mM cholesterol. The cells were washed with KRBH buffer (x2), incubated with KRBH/0.1 % (w/v) BSA (pH 6.4) without or with IF_1_ (final concentration 10 µg/mL), incubated with apoA-I (final concentration 1 mg/mL) for 1 h, then incubated for 10 min with MitoSOX (1 µM, Thermo Fisher) and Hoechst 33342 (1:200 (v/v)). The samples were washed with KRBH and mitochondrial oxidative stress was quantified by confocal laser scanning microscopy as described in Supplementary Methods

Localization of internalized apoA-I in the Ins-1E cells was determined by incubation for 1 h at 37 °C with unlabelled apoA-I (final concentration 0.9 mg/mL) and AF488-labelled apoA-I (final concentration 0.1 mg/mL), then incubation at 37 °C for 10 min with MitoTracker Red CM-H2XRos (Thermo Fisher). The cells were washed with PBS, fixed with 10% (v/v) formalin, washed again with PBS, then imaged with a Zeiss LSM880 microscope in super resolution (Airyscan) mode. Mander’s coefficient for % AF488 overlap with MitoTracker was calculated by comparing the AF488 and MitoTracker channels for the ‘apoA-I’ group or the MitoTracker channel flipped 90°clockwise for the ‘control’ group, using the Just Another Colocalization Plugin (JACoP) in FIJI software (NIH, Bethesda, MD).

### Statistical analyses

Results are presented as the mean±SEM. Statistically significant differences between data sets were identified by one sample Student’s t-test (p values denoted by #), unpaired, two-tailed Student’s t-test (p values denoted by *), and 1-way ANOVA or 2-way ANOVA with Sidak’s Tukey’s multiple comparison test (p values denoted by *) where appropriate. GraphPad Prism (version 9.4, San Diego, CA) was used for all analyses with p<0.05 considered as statistically significant.

## RESULTS

### ApoA-I increases GSIS in non-cholesterol-loaded and cholesterol-loaded Ins-1E cells

Incubation of Ins-1E cells with cholesterol-methyl-β-cyclodextrin in the absence of apoA-I increased intracellular cholesterol levels from 24±1.7 to 41±2.4 µg/mg protein (Fig 1A, p<0.001 vs control), and to 38±2.2 µg/mg protein in the presence of apoA-I (Fig 1A, p<0.01 vs control).

When cholesterol-loaded Ins-1E cells were incubated under basal (2.8 mM glucose) conditions, insulin secretion decreased by 44±4.3%, compared to Ins-1E cells with normal cholesterol levels (Fig 1B, p<0.01). In the presence of apoA-I, insulin secretion increased by 95±8.4% compared to cholesterol-loaded cells incubated without apoA-I (Fig 1B, p<0.01).

The same pattern was observed under high (25 mM) glucose conditions, where GSIS in cholesterol-loaded Ins-1E cells decreased by 60±9.7% compared to control Ins-1E cells (Fig 1C, p<0.01). Incubation with apoA-I restored GSIS in the cholesterol-loaded Ins-1E cells back to that of cells with normal cholesterol levels (Fig 1C, p<0.05 versus cholesterol-loaded cells without apoA-I).

Incubation of non-cholesterol loaded Ins-1E cells with apoA-I under basal (Fig 1B, 2.8 mM) and high (Fig 1C, 25 mM) glucose conditions increased insulin secretion by 80±22% and 350±116%, respectively, relative to cholesterol-loaded Ins-1E cells incubated in the presence of apoA-I (Fig 1B-C, p<0.05 for both). Incubation with apoA-I and cholesterol-loading does not affect Ins-1E protein levels (Supplementary Fig. 1) or cell viability (Supplementary Fig. 2).

**Fig 2.**
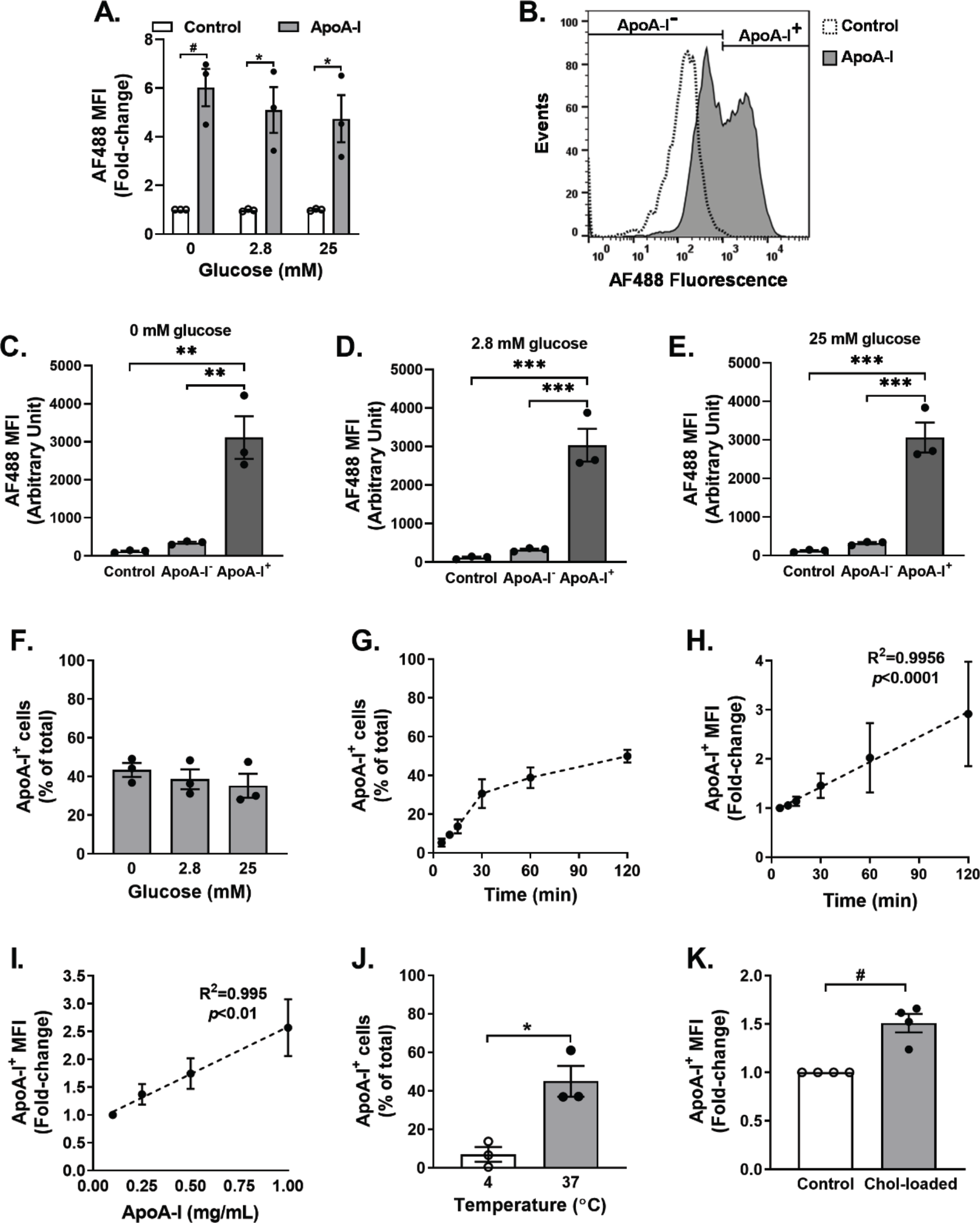
ApoA-I is internalized by a subset of Ins-1E cells in a time-, temperature- and cholesterol-dependent manner. Ins-1E cells were incubated for 1 h at 37 °C in the absence or presence of unlabelled apoA-I (final concentration 0.9 mg/mL), AF488-labelled apoA-I (final concentration 0.1 mg/mL) and 0, 2.8 or 25 mM glucose. AF488 internalization was quantified by flow cytometry. **Panel A** shows mean fluorescence intensity (MFI) of AF488 in cells incubated in the absence (open bars) or presence (closed bars) of 0, 2.8 and 25 mM glucose. Based on AF488 fluorescence relative to control, Ins-1E cells that were incubated with AF488-labelled apoA-I in the absence of glucose were gated as apoA-I^−^ cells or apoA-I^+^ cells (**Panel B**). MFI of AF488 in the control, apoA-I^−^ and apoA-I^+^ subsets of Ins-1E cells incubated with 0 (**Panel C**), 2.8 (**Panel D**) and 25 (**Panel E**) mM glucose are shown. The % Ins-1E cells that internalized AF488-labelled apoA-I in absence or presence of 2.8 and 25 mM glucose is shown in **Panel F**. The % Ins-1E cells that internalized AF488-labelled apoA-I and the AF488 MFI over 120 min are shown in **Panel G** and **Panel H,** respectively. **Panel I** shows the AF488 MFI of apoA-I^+^ Ins-1E cells following incubation for 1 h with unlabelled apoA-I (0.09 – 0.9 mg/mL) and AF488-labelled apoA-I (0.01 – 0.1 mg/mL). **Panel J** shows the % apoA-I^+^ Ins-1E cells after incubation for 1 h at 4 and 37 °C. **Panel K** shows the MFI of apoA-I^+^ cells in control and cholesterol-loaded cells. Values in **Panels A** and **C‒K** represent the mean±SEM of 3-4 independent experiments, with 10,000 events per acquisition. Data in Panels A, J and K were analyzed by 1-sample t-test or unpaired, 2-tailed t-test, as appropriate. Panels C-F were analyzed by 1-way ANOVA, and Tukey’s multiple comparison tests. Simple linear regression was used to calculate the slopes in Panels H and I. *p<0.05, **p<0.01, ***p<0.001, ^#^p<0.05.

### Ins-1E cells selectively internalize apoA-I in a time-, temperature-, concentration- and cholesterol-dependent manner

Although the ability of apoA-I to increase GSIS in Ins-1E cells with normal cholesterol levels has been reported to depend on its internalization^16^, it is not known whether this is also the case for cholesterol-loaded Ins-1E cells. This was addressed by incubating Ins-1E cells with AF488-labelled apoA-I. To ensure that only intracellular fluorescence was quantified, cell surface AF488 fluorescence was quenched at the end of the incubation with an anti-AF488 antibody.

When Ins-1E cells with normal cholesterol levels were incubated with AF488-labelled apoA-I in the presence of 0, 2.8 and 25 mM glucose, fluorescence increased by 502±77%, 420±87% and 374±93%, respectively (Fig 2A, p<0.05 for all). This indicates that apoA-I internalization is independent of glucose concentration.

However, apoA-I was not internalized uniformly. For the incubations in the absence of glucose, cells without (apoA-I^−^ cells) and with internalized apoA-I (apoA-I^+^ cells) were identified (Fig 2B). AF488 fluorescence in the apoA-I^+^ cells was 25-fold higher than in the control cells (3,114±561 versus 121±19 MFI) and 8-fold higher in the apoA-I^−^ cells (3,114±561 versus 347±27 MFI) (Fig 2C, p<0.01 for both). Fluorescence in the apoA-I^+^ Ins-1E cells was also increased relative to control and apoA-I^−^ cells following incubation under basal (2.8 mM, Fig 2D) and high (25 mM, Fig 2E) glucose conditions (p<0.001 for all).

In the presence of 0, 2.8 and 25 mM glucose, the Ins-1E cells with internalized apoA-I comprised 43±3.6%, 39±5.1% and 35±6.2% of the total cells, respectively (Fig 2F). The AF488 MFI in these cells did not change, regardless of the glucose concentration (Supplementary Fig 4A). The % apoA-I^+^ cells increased linearly to 31±7.4% during the first 30 min of incubation and 50±3.2% of the apoA-I was internalized by 2 h (Fig 2G).

ApoA-I internalization, as reflected by AF488 MFI, increased linearly up to 2 h (Fig 2H), and was concentration-dependent (Fig 2I, Supplementary Fig 4B). It was also temperature-dependent, with 6.9±3.8% of the Ins-1E cells containing internalized apoA-I after 1 h at 4 °C compared to 45±8.0% for cells that were incubated for 1 h at 37 °C (Fig 2J, p<0.05). This translates into a 200±6.0% increase in apoA-I internalization at 37 °C relative to 4 °C (Supplementary Fig 4C, p<0.001). ApoA-I internalization was also increased in cholesterol-loaded Ins-1E cells relative to cells that were not cholesterol-loaded (Fig 2K, p<0.05). However, cholesterol loading had no impact on the proportion of cells that internalized apoA-I (47.9±5.6% and 34.9±1.9% for control versus cholesterol-loaded cells, respectively, Supplementary Fig 4D).

### ApoA-I binds to an F_1_-ATPase β-subunit on the Ins-1E cell surface

To identify proteins on the Ins-1E cell surface on which apoA-I internalization is dependent, purified apoA-I was coupled to Dynabeads and added to Ins-1E cells at 4 °C, a temperature at which apoA-I internalization is minimal (Fig 2J, Supplementary Fig 4C). The Ins-1E cell surface proteins that bound to Dynabead-coupled apoA-I were identified by mass spectrometry. The β-subunit of F_1_-ATPase, a mitochondrial protein that is localised on the surface of several cell types, including Ins-1E cells was identified as a main apoA-I binding partner (Supplementary Table 1).^19–22^ The presence of this protein on the Ins-1E cell surface was confirmed by flow cytometry using 3D5, an F_1_-ATPase β-subunit-specific antibody (Fig 3A-B, p<0.05 for isotype control versus 3D5).

**Fig 3.**
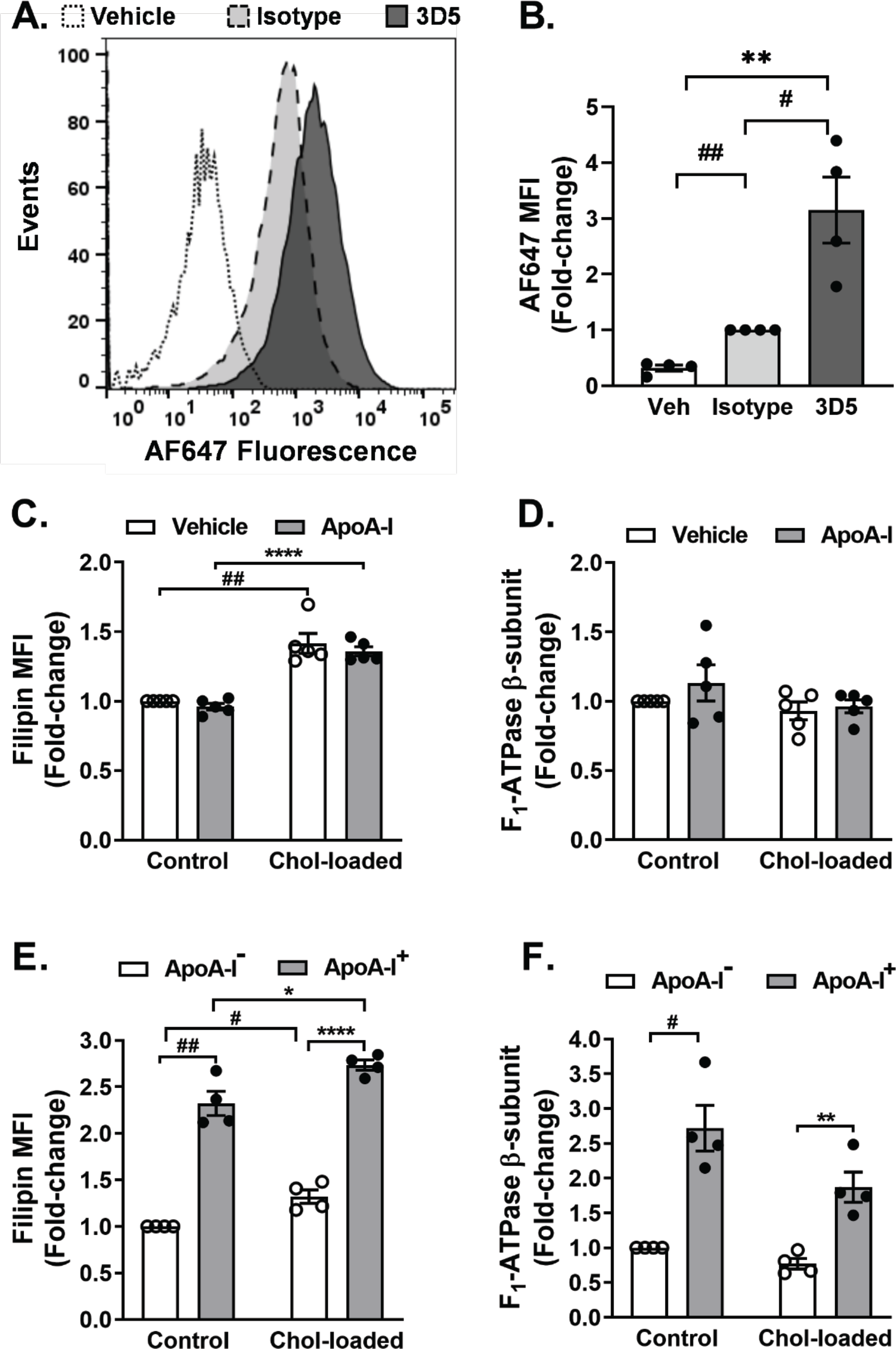
**Cholesterol loading increases apoA-I internalization in Ins-1E cells without altering cell surface F_1_-ATPase β-subunit levels**. Ins-1E cells were incubated in KRBH for 1 h, harvested, fixed with 10% (v/v) formalin and incubated for 1 h at 4 °C with PBS (vehicle), isotype IgG control (1:50, 10 µg/mL) or the 3D5 anti-F_1_-ATPase β-subunit antibody (1:100, 10 µg/mL). The cells were washed with PBS, then incubated for 30 min at 4 °C with an AF647-labelled secondary antibody (1:500). Representative flow cytometry histograms (**Panel A**) and the mean fluorescence intensity (MFI) (**Panel B**) of AF647 are shown. For **Panels C-F**, Ins-1E cells were loaded with 5 mM cholesterol as described in the Legend to Fig 1 then incubated for 1 h in the absence or presence of unlabelled apoA-I (0.9 mg/mL) and AF488-labelled apoA-I (0.1 mg/mL). The cells were harvested, fixed with 10% (v/v) formalin and incubated for 1 h at 4 °C with 3D5 (1:100, 10 µg/mL). After washing with PBS, the cells were incubated for 30 min at 4 °C with an AF647-labelled secondary antibody (1:500), then incubated for 30 min at 4 °C with filipin. MFI of filipin (**Panel C**) and cell surface F_1_-ATPase β-subunit (**Panel D**) in control and apoA-I treated cells are shown. MFI of filipin (**Panel E**) and cell surface F_1_-ATPase β-subunit (**Panel F**) in apoA-I^−^ cells (open bars) and apoA-I^+^ cells (closed bars) are shown. The values in **Panels B–F** represent mean±SEM of 4-5 independent experiments, with 10,000 events per acquisition. Data in Panels B-F were analyzed via 1-sample t-test or unpaired, 2-tailed t-test, as appropriate. *p<0.05, **p<0.01, ****p<0.0001, ^#^p<0.05, ^##^p<0.01

### ApoA-I internalization in Ins-1E cells correlates with cell surface F_1_-ATPase β-subunit, cholesterol and insulin levels

To determine how the cell surface F_1_-ATPase β-subunit regulates apoA-I internalization, unmodified and cholesterol-loaded Ins-1E cells were pre-incubated with AF488-labelled apoA-I, then incubated with the 3D5 antibody. Cellular unesterified cholesterol levels were quantified with filipin.

Cholesterol loading increased filipin fluorescence by 42±7.2% in control Ins-1E cells (Fig 3C, open bars, p<0.01) and by 36±5.3% in the cells that were incubated with apoA-I (Fig 3C, closed bars, p<0.0001). Incubation with apoA-I did not affect filipin fluorescence, irrespective of whether the cells were cholesterol-loaded (Fig 3C). This is consistent with Fig 1A, where cholesterol levels in cholesterol-loaded Ins-1E cells were not altered by incubation with apoA-I. Neither cholesterol loading, nor incubation with apoA-I, altered Ins-1E cell surface F_1_-ATPase β-subunit levels (Fig 3D).

As only a proportion of the Ins-1E cells internalized apoA-I (Fig 2B-E), the impact of this heterogeneity on cell surface F_1_-ATPase β-subunit levels was evaluated. Unmodified and cholesterol-loaded Ins-1E cells were incubated with apoA-I and sorted into apoA-I^+^ and apoA-I^−^ cells. Cholesterol and cell surface F_1_-ATPase β-subunit levels were quantified with filipin and 3D5, respectively. In the absence of cholesterol loading, filipin fluorescence in the apoA-I^+^ Ins-1E cells was 132±13% higher than in apoA-I^−^ Ins-1E cells (Fig 3E, p<0.01). Filipin fluorescence in the cholesterol-loaded apoA-I^+^ Ins-1E cells was 109±10% higher than cholesterol-loaded apoA-I^−^ Ins-1E cells (Fig 3E, p<0.0001). Filipin fluorescence in the cholesterol-loaded apoA-I^+^ Ins-1E cells was 19±7.0% higher than in apoA-I^+^ Ins-1E cells that were not loaded with cholesterol (Fig 3E, closed bars, p<0.05).

In the absence of cholesterol loading, cell surface F_1_-ATPase β-subunit levels in apoA-I^+^ Ins-1E cells were 172±33% higher than in apoA-I^−^ Ins-1E cells (Fig 3F, p<0.05). Cell surface F_1_-ATPase β-subunit levels in cholesterol-loaded cells were 142±9% higher in apoA-I^+^ Ins-1E cells than in apoA-I^−^ Ins-1E cells (Fig 3F, p<0.01

Insulin levels were also quantified in Ins-1E cells following incubation with apoA-I in the absence or presence of 2.8 or 25 mM glucose. Insulin levels in apoA-I^+^ cells were 62±11% (ns), 68±6.6% (p<0.05) and 67±4.7% (p<0.01) lower than in apoA-I^−^ Ins-1E cells in the absence or presence of 2.8 and 25 mM glucose (Fig 4A-B).

**Fig 4.**
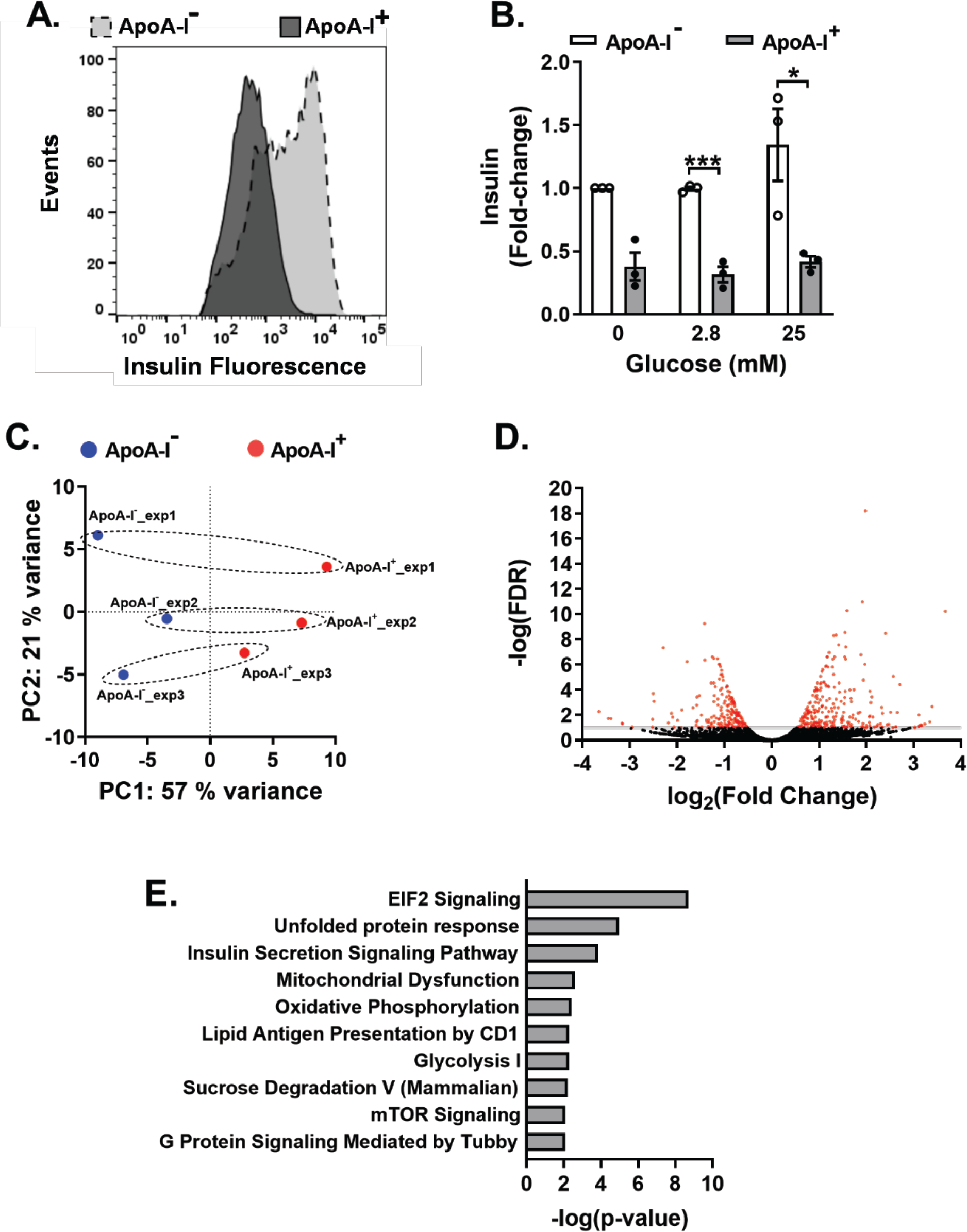
Ins-1E cells that internalize apoA-I have reduced insulin levels and differentially regulated genes. Ins-1E cells were incubated at 37 °C for 1 h with unlabelled apoA-I (0.9 mg/mL) and AF488-labelled apoA-I (0.1 mg/mL) in the absence or presence of glucose (2.8 and 25 mM), then sorted into apoA-I^−^ cells and apoA-I^+^ cells on the basis of AF488 fluorescence. The sorted cells were fixed, permeabilised and incubated for 30 min at 4 °C with an AF647-conjugated anti-insulin antibody. A representative histogram (**Panel A**) and the mean fluorescence intensity (MFI) (**Panel B**) of insulin in the apoA-I^−^ and apoA-I^+^ Ins-1E cells are shown. **Panel C** shows 2D principal component analysis (PCA) plots for apoA-I^+^ and apoA-I^−^ Ins-1E cells from 3 independent experiments. Differentially expressed genes (red) with an FDR of ≤ 0.1 are shown (**Panel D**). The top 10 canonical pathways based on differential gene expression in apoA-I^+^ Ins-1E cells relative to apoA-I^−^ Ins-1E cells were identified by Ingenuity Pathway Analysis (IPA) (**Panel E**). The values in **Panel B** represent the mean±SEM of 3 independent experiments, with 10,000 events per acquisition. Data in Panel B were analyzed via 1-sample t-test or unpaired, 2 tailed t-test, as appropriate. *p<0.05, ***p<0.001.

### Differential gene expression in apoA-I^−^ and apoA-I^+^ Ins-1E cells

Having established that intracellular cholesterol (Fig 3E), cell surface F_1_-ATPase β-subunit (Fig 3F) and total insulin levels (Fig 4A-B) differ significantly in apoA-I^−^ and apoA-I^+^ Ins-1E cells, we next asked whether these differences are also reflected in altered gene expression. ApoA-I^−^ and apoA-I^+^ Ins-1E cells were subjected to RNA-seq and canonical pathways with differentially regulated genes were identified by Ingenuity Pathway Analysis (IPA). Principal component analysis (PCA) showed that gene expression in the apoA-I^−^ and apoA-I^+^ Ins-1E cells clustered separately (Fig 4C), which is consistent with different gene expression profiles in these cell subsets. Out of a total of 6,808 genes, 459 were differentially expressed in apoA-I^−^ and apoA-I^+^ Ins-1E cells, with 258 genes being upregulated, and 201 genes being downregulated in apoA-I^+^ cells relative to apoA-I^−^ Ins-1E cells (FDR ≤ 0.1, Fig 4D). After excluding 121 unidentified differentially expressed genes, the remaining 338 genes were analysed by IPA. The top five pathways that were significantly altered were eukaryotic initiation factor 2 (eIF2) signalling [-log(p-value): 8.69], unfolded protein response (UPR) [-log(p-value): 4.97], insulin secretion signalling [-log(p-value): 3.85], mitochondrial dysfunction [-log(p-value): 2.61] and oxidative phosphorylation [-log(p-value): 2.42] (Fig 4E, Supplementary Table 2). Seventeen genes involved in eukaryotic initiation factor 2 (eIF2) signalling and 7 genes involved in the unfolded protein response (UPR) were upregulated in apoA-I^+^ Ins-1E cells relative to apoA-I^−^ Ins-1E cells (Supplementary Table 2).

Increased mRNA levels of chaperones such as calnexin (*Canx*), calreticulin (*Calr*), UBX domain protein 4 (*Ubxn4*) and X-box binding protein 1 (*Xbp1*) that target proteins for correct folding or degradation in the UPR suggest that the increased ER stress in apoA-I^+^ Ins-1E cells may be caused by increased protein synthesis, and defective protein folding (Supplementary Table 2). Genes associated with insulin signalling and secretion that were upregulated in apoA-I^+^ Ins-1E cells included *Ins1* and *Ins2*. This is in contrast to the result in Fig 4B, which shows that apoA-I^+^ Ins-1E cells have lower insulin levels than apoA-I^−^ Ins-1E cells. Only four genes associated with insulin secretion were downregulated in apoA-I^+^ Ins-1E cells compared to apoA-I^−^ Ins-1E cells. These downregulated genes included translocon subunit gamma (*Sec61g*), which translocates proteins, including preproinsulin, from the cytosol to the ER and ryanodine receptor 1 (*Ryr1*), which regulates β-cell ER Ca^2+^ homeostasis (Supplementary Table 2).

Other differentially expressed genes were associated with oxidative phosphorylation and mitochondrial dysfunction (Supplementary Table 2). Mitochondrial genes encoding for NADH dehydrogenase (Respiratory Complex I), including NADH-ubiquinone oxidoreductase chain 5 protein (*Mt-Nd5*) and NADH-ubiquinone oxidoreductase chain 6 protein (*Mt-Nd6*), were downregulated in apoA-I^+^ Ins-1E cells compared to apoA-I^−^ Ins-1E cells, while other genes such as NADH dehydrogenase (ubiquinone) 1 beta subcomplex subunit 2, mitochondrial (*Ndufb2*) and NADH dehydrogenase (ubiquinone) 1 beta subcomplex subunit 8, mitochondrial (*Ndufb8*) were upregulated. Similarly, mitochondrial genes including cytochrome c oxidase subunit 2 (*Mt-Co2*), a subunit of Respiratory Complex IV and the ATP synthase F_0_ subunit 6 (*Mt-Atp6*) were downregulated in apoA-I^+^ Ins-1E cells compared to apoA-I^−^ Ins-1E cells, while the mitochondrial ATP synthase F_1_ subunit delta (*Atp5f1d*) was upregulated.

### Internalized apoA-I localises to mitochondria in Ins-1E cells

Having established that several genes involved in mitochondrial redox function are differentially regulated in apoA-I^+^ Ins-1E cells, we next asked whether internalized apoA-I localises to the mitochondria.

When Ins-1E cells were incubated with AF488-labelled apoA-I and MitoTracker, then subjected to confocal microscopy, a high degree of co-localisation of internalized AF488-labelled apoA-I with MitoTracker was observed (Fig 5A-C). The Mander’s coefficient of co-localisation was 0.7±0.1 (Fig 5D, p<0.01).

**Fig 5.**
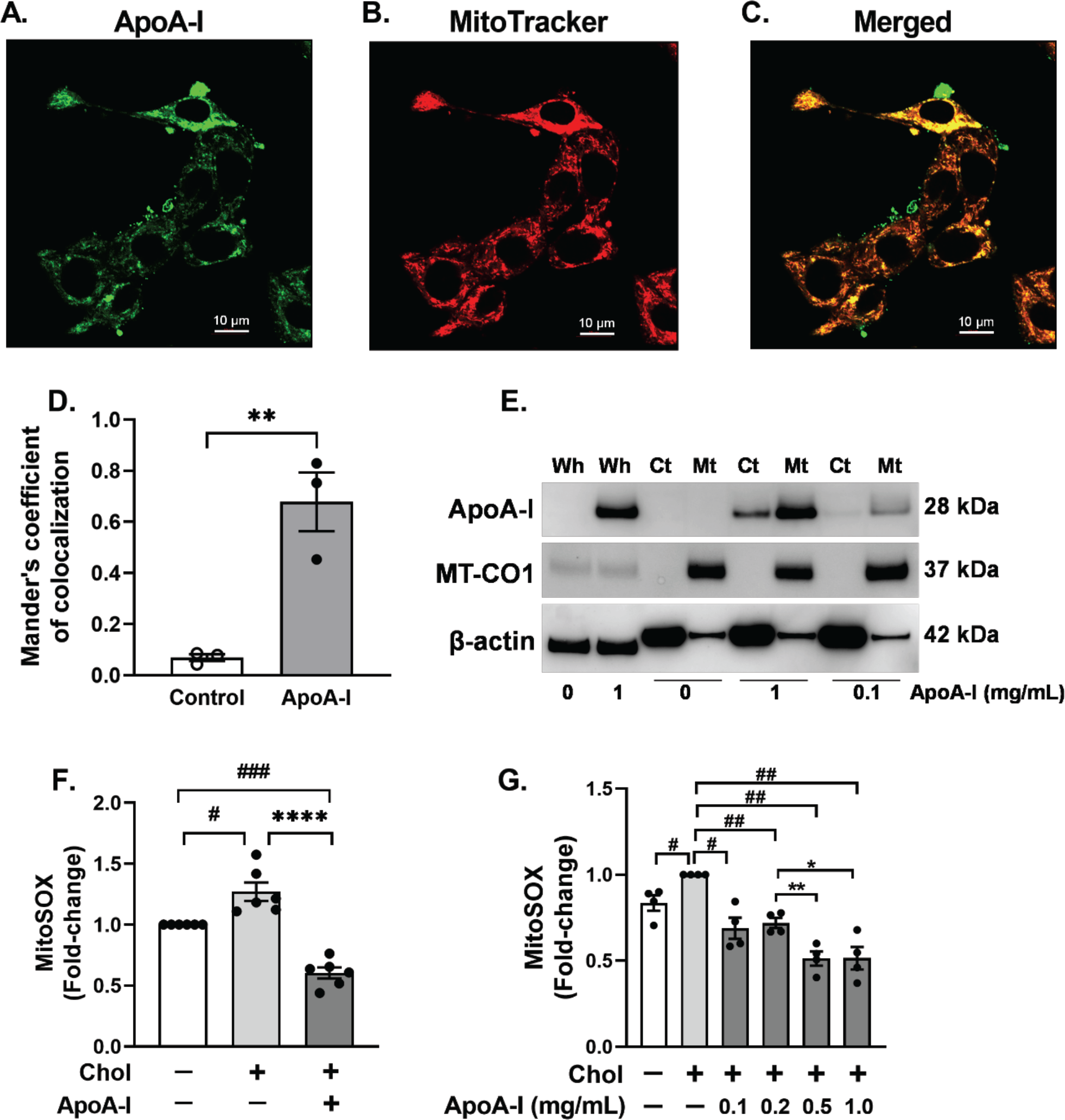
ApoA-I colocalises with mitochondria and reduces cholesterol-induced ROS generation in Ins-1E cells. Ins-1E cells were incubated at 37 °C for 1 h with unlabelled apoA-I (0.9 mg/mL) and AF488-labelled apoA-I (0.1 mg/mL), then incubated for 10 min with MitoTracker. The cells were washed, fixed and AF488 fluorescence was evaluated using a Zeiss LSM880 microscope with super resolution (Airyscan) mode. Fluorescence images of apoA-I (**Panel A**) and MitoTracker (**Panel B**) and a merged image (**Panel C**) are shown. **Panel D** shows the Mander’s coefficient of AF488 overlap with MitoTracker. This result is the mean±SEM of 3 independent experiments with 6-9 images/experiment. **Panel E** shows a representative western blot (from 3 independent experiments) of apoA-I, MT-CO1, and β-actin in whole cell lysates (Wh), and the cytoplasmic (Ct) and mitochondrial (Mt) fractions from Ins-1E cells incubated at 37 °C for 1 h in absence or presence of apoA-I (0.1 or 1 mg/mL). **Panel F** shows MitoSOX fluorescence in control and cholesterol-loaded Ins-1E cells after 1 h of incubation at 37 °C with KRBH or apoA-I (1 mg/mL). **Panel G** shows MitoSOX fluorescence in cholesterol-loaded Ins-1E cells, following incubation at 37 °C for 1 h in the absence or presence of apoA-I (0.1 – 1.0 mg/mL). MitoSOX fluorescence was normalised to cell protein. The values in **Panels F-G** represent the mean±SEM of 4-6 independent experiments, with 6 replicate samples for each independent experiment. Data in Panels D, F and G were analysed via 1-sample t-test or unpaired, 2-tailed t-test, as appropriate. *p<0.05, **p<0.01, ****p<0.0001, ^#^p<0.05, ^##^p<0.01, ^###^p<0.001.

Co-localisation of internalized apoA-I with mitochondria was confirmed by western blotting of whole cells and mitochondrial and cytoplasmic subfractions (Fig 5E). ApoA-I was not detected in the whole cell lysates or mitochondrial or cytoplasmic subfractions in cells that were not incubated with apoA-I (Fig 5E). Following incubation with apoA-I, whole cell lysates and the mitochondrial fraction were apoA-I-enriched relative to the cytoplasmic fraction (Fig 5E). MT-CO1, a mitochondrial-enriched protein, and β-actin (loading control) are also shown (Fig 5E). The different migration of β-actin in the mitochondrial and cytoplasmic fractions likely reflects the use of a proprietary buffer for cellular subfractionation, while RIPA buffer was used for whole cell lysates.

### ApoA-I reduces mitochondrial ROS levels in cholesterol-loaded Ins-1E cells

Elevated cholesterol levels increase mitochondrial ROS production and reduce mitochondrial anti-oxidant enzymes, leading to GSIS suppression.^23, 24^ Conversely, mitochondrial-targeted anti-oxidants reduce oxidative stress and improve β-cell insulin secretion.^25^ To determine if the apoA-I that co-localised with mitochondria in cholesterol-loaded apoA-I^+^ Ins-1E cells reduces mitochondrial ROS generation, unmodified and cholesterol-loaded Ins-1E cells were incubated in the absence or presence of apoA-I. Mitochondrial ROS levels were quantified with MitoSOX.

Cholesterol loading increased mitochondrial ROS formation in the Ins-1E cells by 27±7.6% compared to Ins-1E cells with normal cholesterol levels (Fig 5F, p<0.05). Inclusion of apoA-I in the incubations decreased mitochondrial ROS formation by 51±5.2% (Fig 5F, p<0.001) in a concentration-dependent manner (Fig 5G, p<0.01). However, apoA-I had no effect on mitochondrial oxygen consumption rate (OCR) or basal respiration in Ins-1E cells (Supplementary Fig. 5A-B). This indicates that apoA-I does not affect mitochondrial respiration.

### Reducing ROS production with the mitochondrial antioxidant mitoTEMPO increases insulin secretion in cholesterol-loaded Ins-1E cells

To establish whether decreasing mitochondrial ROS generation was associated with increased insulin secretion, cholesterol-loaded Ins-1E cells were incubated in the absence or presence of mitoTEMPO, a mitochondrial-targeted antioxidant. The results showed that cholesterol loading decreased insulin secretion in Ins-1E cells by 54±6.5% (Supplementary Fig 6, p<0.0001). Incubation of the cholesterol-loaded Ins-1E cells with mitoTEMPO increased insulin secretion by 43±7.7% (Supplementary Fig. 6, p<0.01) compared to cholesterol-loaded Ins-1E cells incubated in the absence of mitoTEMPO.

**Fig 6.**
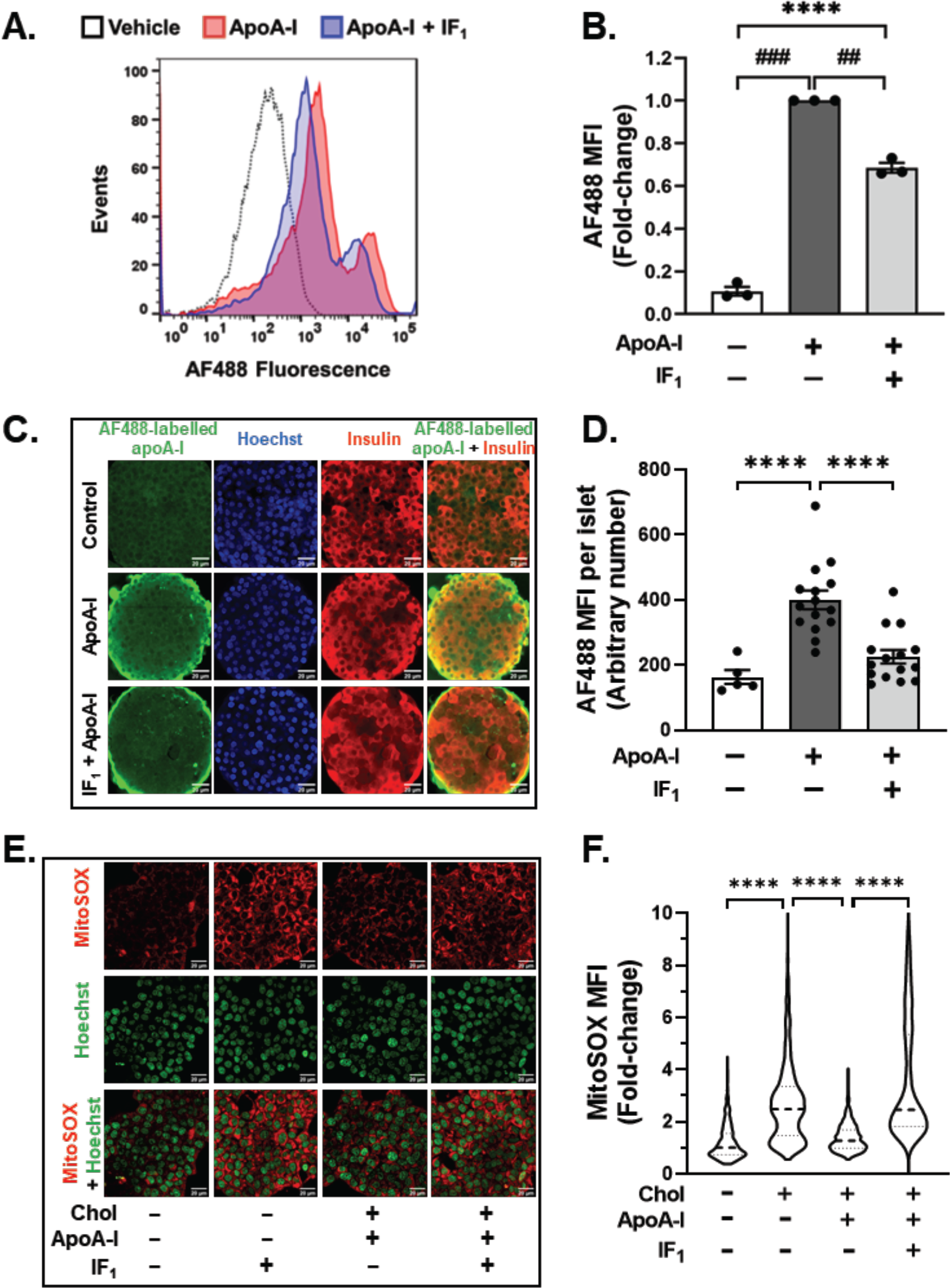
IF_1_ inhibits apoA-I internalization and promotes ROS generation in cholesterol-loaded Ins-1E cells and isolated islets. Ins-1E cells were pre-incubated at 37 °C for 15 min in KRBH without or with IF_1_ (10 µg/mL), then incubated at 37 °C for 30 min with KRBH, 0.1% (w/v) BSA (vehicle) or AF488-labelled apoA-I (0.1 mg/mL) in the presence or absence of IF_1_ (10 µg/mL). Representative histograms (**Panel A**) and MFI (**Panel B**) of AF488 are shown. **Panels C-D** shows isolated islets that were pre-incubated at 37 °C for 15 min in KRBH (pH 6.4) with or without IF_1_ (10 µg/mL), incubated at 37 °C for 2 h with KRBH, 0.1% (w/v) BSA (vehicle) or AF488-labelled apoA-I (0.1 mg/mL), then incubated at 37 °C with Hoechst 33342 (1:200 (v/v)) for 10 min. When the incubations were complete the islets were washed with KRBH, fixed, permeabilised and incubated overnight at 4 °C with an AF647-labelled anti-insulin antibody (1:100). Representative confocal images (**Panel C**) and MFI of AF488 in individual islets (**Panel D**) are shown. For **Panels E-F**, control and cholesterol-loaded cells were incubated for 15 min in KRBH (0.1 % BSA (v/v), pH 6.4) without or with IF_1_ (final concentration: 10 µg/mL). The cells were further incubated for 1 h in the presence or absence of apoA-I (final concentration 1 mg/mL), followed by incubation with MitoSOX (1 µM) and Hoechst 33342 (1:200 dilution (v/v)) for 10 min. Representative confocal images (**Panel E**) and MFI of MitoSOX (**Panel F**) are shown. The **Panel B** represents the mean±SEM of 3-4 independent experiments. **Panel D** represents mean±SEM of 5-15 islets from each of 3 mice. **Panel F** represents the median value of 1185-1474 cells from 3 independent experiments. Data in Panel B was analyzed via 1-sample t-test or unpaired, 2-tailed t-test, as appropriate. Data in Panels D and F were analyzed via 1-way ANOVA, followed by Tukey’s multiple comparison test. ****p<0.0001, ^##^p<0.01, ^###^p<0.001.

### ApoA-I decreases mitochondrial ROS levels in cholesterol-loaded Ins-1E cells and isolated islets in a cell surface F_1_-ATPase β-subunit-dependent manner

We next tested whether inhibiting apoA-I binding to cell surface F_1_-ATPase β-subunit with IF_1_, a mitochondrial ATPase inhibitory factor blocks internalization.^22, 26^ IF_1_ inhibited AF488-labelled apoA-I internalization in Ins-1E cells by 31±2.3 % (Fig 6A-B, p<0.01). This suggests that apoA-I internalization is dependent, at least in part, on binding to the F_1_-ATPase β-subunit.

This result was recapitulated in incubations of isolated mouse islets with AF488-labelled apoA-I and IF_1_. Relative to islets incubated with apoA-I only, the mean fluorescence intensity of internalized AF488-labelled apoA-I decreased by 44±9.0% in the presence of IF_1_ (Fig 6C-D, p<0.0001). The presence of β-cells in the islets was confirmed by staining for insulin (Fig 6C). Merging of the AF488-labelled apoA-I and insulin images showed co-localisation of apoA-I within insulin-positive cells in the islets that were incubated with apoA-I. This was reduced by incubation in the presence of IF_1_ (Fig 6C).

In agreement with the MitoSOX data in Fig 5E and 5F, confocal microscopy further established that the median fluorescence intensity of MitoSOX in cholesterol-loaded Ins-1E cells was increased by 148±1.7% relative to control (Fig 6E-F, p<0.0001). In the presence of apoA-I, MitoSOX fluorescence intensity and mitochondrial ROS generation in cholesterol-loaded Ins-1E cells decreased by 49±1.2% (Fig 6F, p<0.0001). Inclusion of IF_1_ in the incubation increased mitochondrial ROS generation by 92±2.7% compared to cholesterol-loaded cells incubated only with apoA-I (Fig 6F, p<0.0001). Collectively, these results indicate that apoA-I internalization is likely responsible for the observed improvement in mitochondrial function.

## DISCUSSION

Elevated cholesterol levels are associated with a reduced β-cell insulin secretion.^1–3^ This has been attributed, at least in part, to oxidative stress as a result of mitochondrial cholesterol accumulation^6, 7^ and mitochondrial dysfunction.^23^ Depleting cholesterol from cholesterol-loaded ý-cells restores insulin secretion.^2^ ApoA-I also increases insulin secretion in Ins-1E and MIN6 cells but this is independent of cellular cholesterol levels.^12–14^ The present study establishes that apoA-I increases insulin secretion in cholesterol-loaded Ins-1E cells and in isolated mouse islets by binding to an ectopic, cell surface mitochondrial F1-ATPase β-subunit. This facilitates the internalization of apoA-I into the cell where it colocalizes with mitochondria and decreases oxidative stress, leading to increased insulin secretion.

It has been reported previously that apoA-I internalization is necessary for increasing insulin secretion in Ins-1E cells and isolated islets with normal cholesterol levels.^16^ The present study established that this is also the case for Ins-1E cells with elevated cholesterol levels. We also found that apoA-I is internalized in a subset of cholesterol-loaded Ins-1E cells with reduced insulin levels. This is consistent with other reports of the insulin secretory capacity of Ins-1E cells being variable.^27^

The present study further establishes that apoA-I internalization by Ins-1E cells and isolated islets is dependent on binding to an ectopic, cell surface F_1_-ATPase β-subunit. The F_1_-ATPase β-subunit is part of the mitochondrial ATP-producing machinery that includes F_0_F_1_ ATPase (ATP synthase or complex V). The F_1_-ATPase β-subunit is mainly localised on the inner mitochondrial membrane of eukaryotic cells, but it has also been reported on the surface of Ins-1E cells, endothelial cells and hepatocytes.^19, 28^ Although the binding of apoA-I to the F_1_-ATPase β-subunit has been reported to mediate HDL uptake in hepatocytes and vascular endothelial cells via the purinergic receptor P2Y13^22, 29^, this receptor was not identified as an HDL binding partner by mass spectrometry and therefore seems not to have a role in apoA-I internalization in Ins-1E cells.

The binding of apoA-I to the F_1_-ATPase β-subunit on the hepatocyte surface increases ATP hydrolysis and stimulates a purinergic G protein-coupled P2Y receptor signalling cascade that culminates in HDL endocytosis. It also increases survival and proliferation of vascular endothelial cells and drives the transcytosis of apoA-I and HDLs across vascular endothelial cells into the subendothelial space.^26, 29, 30^ How the F_1_-ATPase β-subunit translocates from mitochondria to the cell surface is unclear but appears to depend on brefeldin A in neural cells.^31^ Whether this is also the case in other cell types is not known. Mitochondria-derived vesicles can also transport proteins from the mitochondria towards peroxisomes and lysosomes, and possibly towards the plasma membrane.^32, 33^ Whether these mechanisms are responsible for the translocation of the F_1_-ATPase β-subunit from mitochondria to the Ins-1E cell surface remains to be determined.

One of the most important observations from the present study is preferential internalization of apoA-I by Ins-1E cells with elevated cholesterol levels. This is consistent with what has been reported for apoA-I internalization in cerebral microvascular endothelial cells via an endocytic process that is inhibited when cell cholesterol levels are reduced with methyl β-cyclodextrin.^34^ However, in contrast to a previous study where cholesterol loading increased translocation of the F_1_-ATPase β-subunit to the surface of vascular endothelial cells,^35^ we found that cholesterol loading had no effect on cell surface F_1_-ATPase β-subunit levels in Ins-1E cells.

Heterogeneity in the insulin secretory capacity of mouse β-cells and human islets has been documented previously.^36–38^ This is consistent with the elevated cholesterol and cell surface F_1_-ATPase β-subunit levels, lower insulin levels and functional heterogeneity of the Ins-1E cells with internalized apoA-I in the present study. Additional evidence of functional heterogeneity of Ins-1E cells comes from the observation that genes involved in ER stress and the UPR are upregulated in apoA-I^+^ Ins-1E cells. ER stress also impairs the ABCA1-mediated efflux of cholesterol from hepatocytes.^39^ This may explain why cholesterol levels were increased in the apoA-I^+^ Ins-1E cells. However, it is unclear if this is a cause or consequence of ER stress.

Reduced expression of *Sec61g* and *Ryr1* that respectively encode for the transport proteins Sec61 subunit gamma and ryanodine receptor 1 in the ER membrane further suggest that ER-associated oxidative folding of insulin and conversion of preproinsulin to proinsulin is decreased in apoA-I^+^ Ins-1E cells.^40–42^ This may explain why apoA-I^+^ Ins-1E cells have lower insulin levels than apoA-I^−^ Ins-1E cells despite having increased *Ins1* and *Ins2* expression (Fig 4A-B). It is also consistent with decreased expression of mitochondrial genes involved in oxidative phosphorylation and function that produce ATP and drive insulin secretion in apoA-I^+^ Ins-1E cells (Supplementary Table 2).

Heterogeneity in UPR activation and ER oxidative stress have been reported in isolated human islets.^43^ This may be caused by β-cells cycling from periods of (i) high insulin synthesis and low UPR to (ii) ER stress, increased UPR and low insulin synthesis to (iii) resolved ER stress and low insulin synthesis.^44^ It is possible that the Ins-1E cell heterogeneity in the present study reflects the transitioning of Ins-1E cells across similar dynamic states, rather than stable subtypes. It is also possible that the Ins-1E heterogeneity in the present study reflects ‘hub’ Ins-1E cells that internalize apoA-I and are highly metabolic, transcriptionally immature and have low insulin levels.^45^ The internalization of apoA-I by ‘hubs’ could modulate function and alter the insulin secretory capacity of cells that do not internalize apoA-I. ý-cells that are defined as “hubs” underpin the responses of islets to changes in glucose levels, making them essential for GSIS.^45^ Moreover, as the viability of ý-cell “hubs” is diminished by glucotoxicity and glucolipotoxicity, it follows that these cells may play a fundamentally important role in diabetes prevention.^45^

The current results indicate that cholesterol enrichment exacerbates ER and oxidative stress and reduces insulin secretory capacity in Ins-1E cells. The antioxidant functions of apoA-I have been well characterised^46^ and have been attributed to reduced levels of ROS and increased expression of antioxidant enzymes including superoxide dismutase and glutathione peroxidase.^47^ The present study indicates that apoA-I internalization may increase the insulin secretory capacity of Ins-1E cells with elevated cholesterol levels by inhibiting mitochondrial ROS production and restoring mitochondrial redox balance. Further studies are needed to determine whether comparable β-cell subpopulations are present in human islets, and whether they are potential targets for pharmacological inhibition of oxidation.

A limitation of the present study is that relative to its level in normal human plasma, the concentration of lipid-free apoA-I used in the GSIS experiments could be considered as supraphysiological. However, it is important to note that lipid-free and lipid-associated apoA-I (as a constituent of reconstituted HDLs) both increase GSIS equally effectively.^12^ This is consistent with the capacity of apoA-I to increase insulin secretion being independent of whether it is lipid-free or lipid-associated. It therefore follows the apoA-I concentration to which the Ins-1E cells were exposed in the GSIS experiments is physiologically relevant.

In conclusion, this study establishes that apoA-I restores insulin secretion in β-cells with disrupted cholesterol homeostasis by binding to an ectopic F1-ATPase β-subunit, internalizing into the cells and localizing to mitochondria where it decreases oxidative stress and enhances insulin secretion in response to glucose. Whether the observed increase in insulin secretion can be attributed to nuclear exclusion of Pdx1, as reported previously, or whether it reflects improved processing of proinsulin to insulin and increased expression of the prohormone convertase 1/3 (PC1/3) remains to be determined.^13, 16^ These insights indicate that increasing apoA-I levels may benefit hypercholesterolemic patients with impaired β-cell function.

## ACKNOWLEDGEMENTS

BM was the recipient of a University International Postgraduate Award, supported by UNSW Sydney and Commonwealth of Australia. S-YC is supported by a UNSW Scientia PhD Scholarship.

## SUPPLEMENTARY MATERIALS

Supplemental Methods

Supplementary Table 1-2

Supplementary Figures 1-5

## DATA AVAILABLITY

RNA-seq datasets have been deposited into ENA (https://www.ebi.ac.uk/ena/browser/) with a primary accession number PRJEB50345.

Mass spectrometry data can be accessed at https://datadryad.org/stash with the link (https://datadryad.org/stash/share/zrWZc9Mk1tVpOcGZdH9Dz6KngftJcQciplpKtC6EGos)

